## Supplemental_material for "Applications of Machine Learning in Decision Analysis for Dose Management for Dofetilide"

### Supplemental Methods

Packages used for analysis include the following:

Unsupervised Learning:

- *sklearn.decomposition.PCA* for principal component analysis
- *sklearn.cluster.KMeans* for K-means clustering with 8 clusters.

Supervised Learning:

- *sklearn.linear_model.LogisticRegression* for L1 regularized logistic regression algorithm with ‘liblinear’ solver
- *sklearn.ensemble.RandomForestClassifier* for Random Forest classification with 500 estimators, and maximum leaf nodes of 20
- *sklearn.ensemble.AdaBoostClassifier* for Boosted decision tree classification, combined with
- *sklearn.tree.DecisionTreeClassifier* with 200 estimators, SAMME.R algorithm, and learning rate of 0.5
- *sklearn.svm.SVC* for support vector machine classification with radial basis function kernel, gamma of 5, and C-value of 0.001
- *sklearn.neighbors.KNeighborsClassifier* for K-nearest neighbors classification with 1 and 10 nearest neighbors

| Parameter | No change | Dose change |
|---|---|---|
| age_years | 7.56E+185 | 1028.198506 |
| bmi | 3.25E+185 | -8.365964 |
| hr_ | 5.38E+185 | -1161.59328 |
| qtc_cpu_based_ | 4.57E+185 | -941.241576 |
| tte_lvef | 6.79E+185 | -848.112759 |
| potassium | 5.30E+185 | 276.983848 |
| magnesium | 4.46E+185 | -170.593196 |
| creatinine | 2.48E+185 | 721.813547 |
| betablocker | -1.01E+185 | 1995.601825 |
| ccb | -2.79E+185 | -640.978418 |
| qrs_duration_ | 3.95E+184 | -144.240687 |
| sex | 3.99E+185 | -189.105307 |
| indication | 3.35E+74 | -0.315224 |
| SR_ | 3.07E+185 | 844.658958 |
| ppm | 6.01E+185 | 2169.177153 |
| CV_ | 1.44E+185 | 1733.051941 |
| dose_ | 9.93E+185 | -3406.121237 |
| htn | 2.95E+182 | 102.409849 |
| icd | 9.86E+185 | -457.81708 |
| afib | -7.61E+176 | 279.38327 |
| vt | 9.87E+185 | -861.85385 |
| dm | 6.01E+185 | -277.424116 |
| cad | 5.87E+185 | -1707.796601 |
| chf | 9.86E+185 | -2100.37959 |
| dose | 3.23E+185 | 373.687832 |

**Supplemental Table 1. Weights for most accurate predictive model.** Alpha = 0.1, gamma = 1.0; accuracy = 0.939.

**Supplemental Figure 1.** Accuracy for linear policy approximation applied to models created from various values of alpha and gamma parameters.

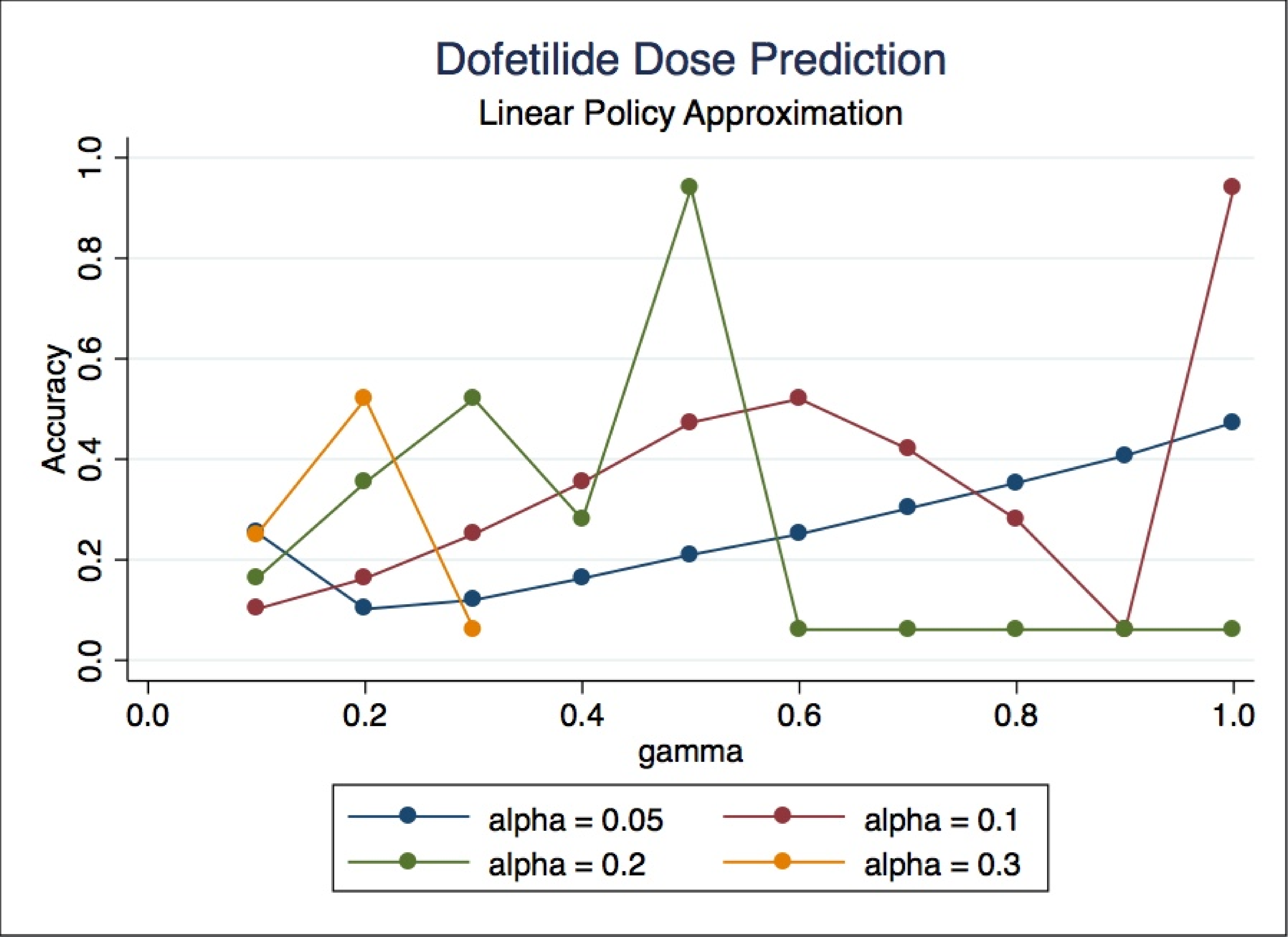
Dofetilide Dose Prediction
Linear Policy Approximation
Accuracy
0.0
0.2
0.4
0.6
0.8
1.0
gamma
0.0
0.2
0.4
0.6
0.8
1.0
alpha = 0.05
alpha = 0.1
alpha = 0.2
alpha = 0.3
